# Peripheral alcohol metabolism dictates ethanol consumption and drinking microstructure in mice

**DOI:** 10.1101/2025.01.09.632203

**Authors:** Bryan Mackowiak, David L. Haggerty, Taylor Lehner, Yu-Hong Lin, Yaojie Fu, Hongkun Lu, Robert J. Pawlosky, Tianyi Ren, Wonhyo Seo, Dechun Feng, Li Zhang, David M. Lovinger, Bin Gao

## Abstract

**Background:** Ethanol metabolism is intimately linked with the physiological and behavioral aspects of ethanol consumption. Ethanol is mainly oxidized by alcohol dehydrogenase (ADH) to acetaldehyde and further to acetate via aldehyde dehydrogenases (ALDHs). Understanding how ethanol and its metabolites work together to initiate and drive continued ethanol consumption is crucial for identifying interventions for alcohol use disorder (AUD). Therefore, the goal of our study was to determine how ADH1, which is mainly peripherally-expressed and metabolizes >90% of ingested ethanol, modulates ethanol metabolite distribution and downstream behaviors.

**Methods:** Ethanol consumption in drinking-in-the-dark (DID) and two-bottle choice (2BC) drinking paradigms, ethanol metabolite concentrations, and lickometry were assessed after ADH1 inhibition and/or in *Adh1*-knockout (*Adh1* KO) mice.

**Results:** We found that *Adh1* KO mice of both sexes exhibited decreased ethanol consumption and preference compared to wild-type (WT) mice in DID and 2BC. ADH1 inhibitor fomepizole (4-MP) also significantly decreased normal and sweetened ethanol consumption in DID studies. Measurement of ethanol and its metabolites revealed that ethanol was increased at 1h but not 15 min, peripheral acetaldehyde was slightly decreased at both time points, and ethanol-induced increases in acetate were abolished after ethanol administration in *Adh1* KO mice compared to controls. Similarly, ethanol accumulation as a function of consumption was 2-fold higher in *Adh1* KO or 4-MP treated mice compared to controls. We then used lickometry to determine how this perturbation in ethanol metabolism affects drinking microstructure. *Adh1* KO mice consume most of their ethanol in the first 30 min like WT mice but display altered temporal shifts in drinking behaviors and do not form normal bout structures, resulting in lower ethanol consumption.

**Conclusions:** Our study demonstrates that ADH1-mediated ethanol metabolism is a key determinant of ethanol consumption, highlighting a fundamental knowledge gap around how ethanol and its metabolites drive ethanol consumption.

## Introduction

Ethanol metabolism is a key determinant of how individuals perceive and experience ethanol (Cederbaum, 2012, Israel et al., 2015). Upon ingestion, ethanol is first metabolized to acetaldehyde mainly by alcohol dehydrogenase 1 (ADH1), with catalase and cytochrome P450 2E1 (CYP2E1) playing minor roles. However, *Adh1* and its human homologs are not expressed in the brain, where catalase is the major enzyme involved in ethanol oxidation (Galter et al., 2003, Cederbaum, 2012). Acetaldehyde is mainly converted to acetate by aldehyde dehydrogenase 2 (ALDH2) throughout the body, with gut and liver ALDH2 clearing the majority of systemic acetaldehyde (Cederbaum, 2012, Fu et al., 2024, Guillot et al., 2019). While ethanol and acetate are well-known to pass the blood-brain barrier (BBB) (Jiang et al., 2013), the ability of AcH to cross the BBB at low concentrations is controversial, though our recent study suggests that increased peripheral AcH can increase brain AcH concentrations (Jamal et al., 2016, Fu et al., 2024) **(Fig. 1)**. Over 90% of all ethanol metabolism occurs in the periphery, but how peripheral ethanol metabolism contributes to the acquisition and maintenance of ethanol consumption is unclear (Cederbaum, 2012).

**Figure 1:**
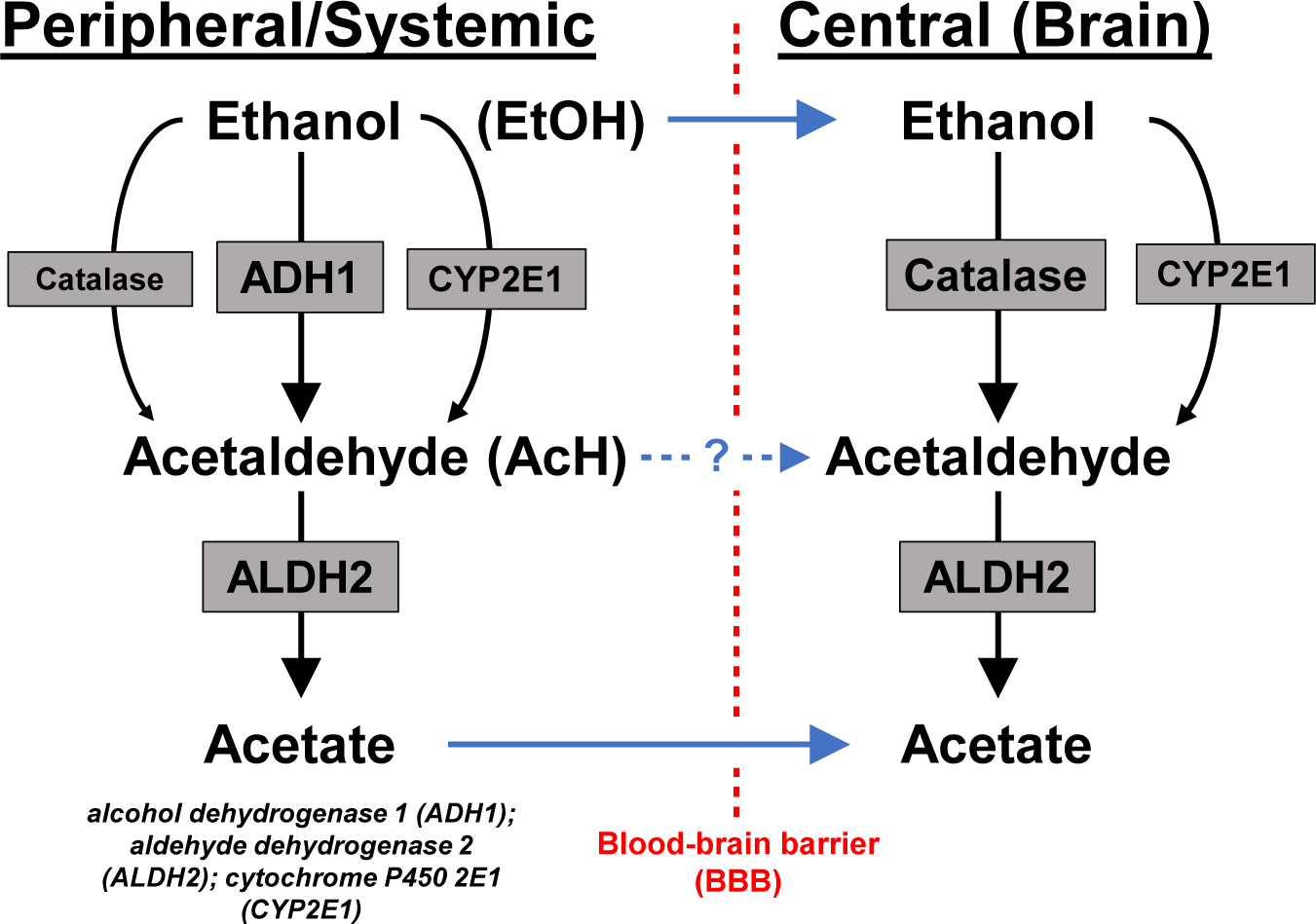
Schematic of alcohol metabolism.

Ethanol, AcH, and acetate all also contribute to the behavioral effects associated with ethanol intoxication (Israel et al., 2015, Karahanian et al., 2011). Ethanol itself alters signaling in many different brain regions, exerting its effects through a variety of direct and indirect molecular targets including direct modulation of γ-aminobutyric acid (GABA) and N-Methyl-D-Aspartate (NMDA) receptors (Koob, 2004, Davis and Wu, 2001, Abrahao et al., 2017). While accumulation of toxic AcH after ethanol consumption causes the aversive flushing reaction along with nausea and palpitations, AcH is also important in the locomotor, motivational, and reinforcing effects of ethanol (Peana et al., 2010, Peana et al., 2008, Peana and Acquas, 2013). In addition, acetate influences ethanol-induced behavior in several ways, including downstream conversion to GABA, activation of NMDA receptors (NMDARs), and modification of histone acetylation (Mews et al., 2019, Jin et al., 2021, Pardo et al., 2013, Chapp et al., 2023, Chapp et al., 2024, Chapp et al., 2014).

As ethanol metabolism is the major factor determining ethanol pharmacokinetics, changes in ethanol metabolism significantly affect neural responses to ethanol and thus the resultant drinking behaviors. This process is primarily driven by alterations in peripheral and central levels of ethanol and its metabolites (Lehner et al., 2024, Haass-Koffler et al., 2017). For instance, genetic polymorphisms in *ADH1B*, *ADH1C*, and *ALDH2* are highly associated with ethanol consumption and the risk of alcohol use disorder (AUD) (Zhou et al., 2022, Luczak et al., 2009, Kimura and Higuchi, 2011). These polymorphisms either increase ADH1 activity or decrease ALDH2 activity, both leading to increased levels of aversive acetaldehyde and initiation of the flushing reaction after ethanol consumption, which decreases the risk of AUD (Kimura and Higuchi, 2011, Edenberg, 2007). To date, most research has focused on the effect of the *ALDH2*2* polymorphism and ALDH2 inhibition on drinking behavior and AUD development. Disulfiram, an ALDH inhibitor, was the first drug approved for AUD treatment, and several second-generation therapies targeting ALDH2 have been developed or are in continued development (HUGHES and COOK, 1997, Lanz et al., 2023, O’Malley et al., 2020, Arolfo et al., 2009, Zhang X, 2022).

While increases in acetaldehyde by ADH1 activation or ALDH2 inhibition clearly decrease ethanol consumption, many other modulations of alcohol metabolism exert differential effects on drinking behavior. Gastric bypass surgery decreases the first-pass metabolism of ethanol in the stomach, leading to a doubling of blood alcohol concentration per drink which contributes to the increased risk of AUD in these patients (Woodard et al., 2011, Acevedo et al., 2020, Blackburn et al., 2017). Chemical modulators of ethanol pharmacokinetics also alter drinking behavior in preclinical models, often in conflicting ways (Lehner et al., 2024, Haass-Koffler et al., 2017). The ALDH2 activator ALDA-1, which decreases AcH accumulation, reduces the acquisition of ethanol intake and chronic ethanol consumption in ethanol-preferring rats, without altering sucrose preference (Rivera-Meza et al., 2019). Metadoxine, a medication known to increase overall ethanol clearance, has been associated with decreased ethanol consumption in two preliminary clinical studies for AUD patients (Guerrini et al., 2006, Leggio et al., 2011, Shpilenya et al., 2002). On the other hand, inhibition of ADH1-dependent ethanol metabolism by the clinically approved competitive inhibitor fomepizole (4-methylpyrazole; 4-MP) was shown to decrease ethanol self-administration in alcohol-preferring rats (Peana et al., 2017) and ethanol-induced conditioned place preference in Wistar rats (Peana et al., 2008). In addition, decreasing ADH activity via immunization against ADH epitopes can decrease ethanol consumption in alcohol-dependent rats (PSHEZHETSKY et al., 1993).

Exactly how alterations in ethanol metabolites specifically translate to measurable differences in ethanol-related behaviors also remains obscure. Monitoring ethanol intake and the associated behaviors has proven a powerful tool to elucidate the temporal dynamics of how rodents consume ethanol. This practice, lickometry, has also unlocked key insights to how AUD develops. Recent studies have used lickometry and other measures of drinking rate across time to show that increased rates of ethanol self-administration ethanol early in access paradigms (*i.e.* “frontloading”) leads to higher risk of developing AUD, and can also be used as a general measure of reward-related behavior (Ardinger et al., 2022, Petersen et al., 2023, Sloan et al., 2020, Carpenter et al., 2019). Further, there have been challenges in determining how ethanol metabolism and tolerance modulate frontloading behaviors. While some studies have suggested frontloading cannot be solely driven by metabolic tolerance, these studies rely on observations from rodents that are bred to prefer alcohol, which often carry multiple mutations in alcohol metabolizing enzymes (Ardinger et al., 2022, Bice et al., 2011). Thus, further work to untangle how ethanol metabolism alters the propensity for intense ethanol drinking would have large implications in understanding how AUD develops.

Direct genetic manipulation of ethanol metabolizing enzymes represents a powerful tool to investigate how ethanol and its metabolites contribute to complex drinking behaviors. Our laboratory and others have successfully used these techniques to unravel how peripheral and central acetaldehyde control ethanol consumption (ISRAEL et al., 2013, Fu et al., 2024, Jin et al., 2021). However, it is not clear how peripheral metabolism of ethanol modulates brain concentrations of ethanol and its metabolites and the resultant behaviors. ADH1 is mainly peripherally-expressed and metabolizes >90% of ingested ethanol, which is principally responsible for peripheral ethanol metabolism (Galter et al., 2003, Cederbaum, 2012). Thus, we used *Adh1* knockout (KO) mice to better understand how peripheral ethanol metabolism shapes drinking behavior.

## Materials and Methods

### Mice

C57BL/6N mice were purchased from Charles River Laboratory and bred in the NIAAA animal facility. *Adh1* knockout (KO) mice were kindly provided by Dr. Duester (Burnham Institute, La Jolla, CA) (Deltour et al., 1999), and backcrossed to a C57BL/6N background for more than 10 generations. *Adh1* KO mice were genotyped as described previously (Anvret et al., 2012, Mackowiak et al., 2022). All mouse experiments described in the current paper were reviewed and approved by the National Institute on Alcohol Abuse and Alcoholism Animal Care and Use Committee in protocols LLD-BG-1 and/or LIN-DL-1.

### Ethanol Consumption Experiments

Male or female mice between 2-8 months old were subjected to several different ethanol consumption protocols essentially as described previously with minor modifications (Guillot et al., 2019, Richardson et al., 2023). **Drinking-in-the-Dark (DID):** Mice were acclimated in single housing for 7 days prior to the start of DID with free access to food and water. Three hours after the start of the dark cycle, water bottles were switched with straight-sipper glass bottles containing 20% ethanol (vol/vol) for 2h on day 1-3 and 4h on day 4 unless otherwise noted. To measure blood and tissue levels of ethanol and its metabolites after DID, the protocol was altered so that on day 4, ethanol bottle switches began 2.5h after the dark cycle started and were staggered every 5min. After 2h of DID, mice were sacrificed and tissues harvested for ethanol and metabolite measurement.

### Modified DID

Mice were subjected to a shortened DID protocol recently published (Richardson et al., 2023) with minor modifications. Mice were singly-housed and placed in a reverse light cycle room for 7 days before experiments started with free access to food and water. Water bottles were switched with either 20% ethanol (vol/vol), sweetened ethanol (20% ethanol + 0.3% saccharin), or 0.1% saccharin 3h after the beginning of the dark cycle on Day 1 for 2h and Day 2 for 4h. Mice were acclimated to ethanol consumption for at least 1 session prior to treatment testing. Day 1 provided a baseline for drinking, reacclimated mice to drinking, and ensured that there was no treatment carry-over from the previous cycle, while day 2 was used to test treatments. Treatments were given 0.5-1h prior to the start of DID. **Two-Bottle Choice (2BC):** Mice were switched to 2-bottle cages and acclimated with free access to two water bottles for ∼1 week prior to initiation of drinking experiments as described previously (Fu et al., 2024). At the start of the experiment, bottles were replaced with two glass tubes fitted with neoprene stoppers and straight stainless-steel sippers containing either water or ethanol (3-20%). Each different ethanol concentration was held for 4 days before increasing to the next concentration. Bottle placement was switched each day and bottle weights were measured at least every 2 days.

### Western Blot and Protein Activity

Western blot and ALDH2 activity assay were performed as described previously (Fu et al., 2024). Briefly, liver tissue was homogenized, prepared, and run according to the ALDH2 activity measurement kit (Abcam, Waltham, MA) instructions. Leftover protein was diluted in RIPA buffer with protease inhibitors (Santa Cruz Biotechnology, Dallas, TX), loading buffer containing beta-mercaptoethanol was added, and samples were boiled at 95⁰C for 10min. Samples were run on a 4-12% Bis-Tris Gel (BioRad, Hercules, CA), transferred to nitrocellulose membrane (BioRad, Hercules, CA), blocked with 5% milk, incubated overnight with primary antibodies for ALDH2 (Abcam, Cat#: ab194587, 1:1000), CYP2E1 (Millipore-Sigma, 1:1000), and β-actin (Sigma-Aldrich, Cat#: A1978), washed with phosphate buffered saline with 0.1% Tween 20 (PBST), incubated with horseradish peroxidase secondary antibodies at room temperature for 1h, washed with PBST, and visualized with SuperSignal™ Western Blot Substrate Pico or Femto.

### Rotarod

Rotarod was conducted as described previously (Fu et al., 2024). Briefly, mice were trained on an accelerating rotarod paradigm (0-300s, 4-40rpm) for at least 3 sessions and were included in experiments if they were able to stay on the rotarod for >250s. On the day of the experiment, the baseline was measured for each mouse before receiving ethanol and rotarod performance was measured at defined timepoints afterwards. Mice were given ethanol via intraperitoneal (IP) injection (1 g/kg), gavage (1 g/kg), or IP injection (2 g/kg) in that order with at least 1 day rest between sessions.

### Ethanol and AcH measurement

Ethanol and AcH were measured via headspace gas-chromatography mass spectrometry (GC-MS) as described previously (Fu et al., 2024, Jin et al., 2021, Pawlosky et al., 2010). Briefly, mice were deeply anesthetized with isoflurane before blood collection from the orbital sinus and subsequent cervical dislocation. The left lobe of the liver and cerebellar cortex tissue were quickly harvested on ice before snap freezing in liquid nitrogen. Whole blood (25µL, ethanol and AcH), serum (25µL, acetate), liver (10-30mg), or cerebellar cortex (10-30mg) were placed into screw-top vials containing ceramic beads for further processing. Perchloric acid (250µL, 0.6N) containing deuterated ethanol and AcH internal standards was added to each tube, vortexed (45s for blood) or bead homogenized at 4⁰C using a Bertin Precellys Evolution, and spun down (12,000g, 15min, 4⁰C). Supernatant (200µL) was transferred to 20mL glass headspace vials for analysis. Sample analysis was conducted on an Agilent Headspace GC-MS system (GC: 7890B, MS: 5977B, Headspace: 7697A) using helium as the carrier gas and methane as the reaction gas. The headspace vials were equilibrated at 70⁰C for 10 min with shaking before a 1 mL injection. Mass/charge ratios of ethanol (47.1), deuterated ethanol (52.1), acetaldehyde (45.1), and deuterated acetaldehyde (49.1) were continuously monitored, and peaks were quantified using MassHunter Quantitative Analysis software (Agilent, v10.2).

### Acetate Measurement

Acetate measurement was conducted via GC-MS as described previously (Pawlosky et al., 2010). Acetate was analyzed as its tertiary butyl dimethylsilyl ether derivative (TMS) using GC-MS in the electron impact mode and quantified using the ^2^H_3_-acetate. **Perchloric acid (PCA) extraction.** To 50 µl of frozen sera 50 µl of a 3.6% perchloric acid solution and 5 µl (32 µM solution) of a sodium ^2^H_3_-acetate (Sigma Aldrich Chemical Co., St. Louis Mo.) was added in 100ul polypropylene conical snap-capped tubes. Samples were vortexed for one minute and neutralized with 5 µl KHCO_3_ (3 M). Approximately 30 mg of frozen brain tissue was added to 100 µl of a 3.6% perchloric acid solution and 10 µl (32 µM solution) of a sodium ^2^H_3_-acetate in 2 ml polypropylene screw capped tubes with 10-15 small glass beads. Samples were shaken on Mini Bead Beater (Biopsec Products, Bartlesville, OK) for 30 seconds (2x) until samples were homogeneous. Samples were placed on ice until centrifugation at 4°C in a Sorvall benchtop centrifuge at a speed of 10.7 × g for 2 min. Eighty microliters of the upper layer was pipetted into clean tubes and neutralized with 9 μl 3M KHCO_3_. **Sample derivatization.** Ten microliters of the sample extracts were evaporated under a stream of nitrogen to dryness in 1.5 mL sylinized screw capped vials. Samples were immediately reacted with 5μl of the sylilating reagent, N-methyl-N-(tri-methylsilyl) trifluoroacetamide (MTBSTFA) with 1% tert-butyldimethylchlorosilane (TBDMCS) reagent and the bis-trimethylfluoro methyl silyl (BSTFA) (Pierce Chemical Co., Rockford, IL) in 15μl of acetonitrile and heated to 60°C for 5 min. **GC-MS determinations.** Samples were analyzed on an Agilent 5973 quadrupole GC-MS (Agilent, Wilmington, DE). One µl of the sample solutions was injected onto a 250μm × 30m capillary DB-1 (Agilent) column in the split injection mode (100:1) using helium as the carrier gas. The injector temperature was set at 250°C and the transfer line at 280°C. The GC oven temperature was programmed from 40 to 325°C at 15°C/min. The mass spectrometer was operated in the electron impact mode (70 eV) and the quadrupole mass analyzer scanned for ions which corresponded to a loss of 15 mass units (-CH_3_) from the molecular ion of the acetate derivative (m/z 117) and its corresponding ^2^H-labeled internal standard (m/z 120) using selected ion monitoring mode. The m/z 117/120 ratio was used to quantify the quantity of acetate in each sample.

### Lickometry Experiments

During modified DID as previously described, lickometry was assessed using custom-built lickometers that used touch-based measures of capacitance to measure bottle contacts with millisecond precision. Briefly, a Feather M0 Adalogger (Adafruit, NY) with an onboard SD card captured filtered capacitive touch data from a MPR121 breakout board (Adafruit, NY) such that when current readings where 12 bits higher or lower than previous readings, a touch or an untouched was registered, respectively. A DS3231 Precision RTC (Adafruit, NY) was used to align those touch and untouched events to date-time values with millisecond precision. A custom-built python script was used to align, filter, and calculate licks, lick durations, bouts, licks per bout, and lick durations per bout using inclusion criteria as previously cited for alcohol drinking experiments (Petersen et al., 2023, Brown et al., 2023). Licking behavior had to have at least 3 licks within 1 second of each other to initiate a bout, and a bout was terminated when there was no licking behavior registered for at least 3 seconds after the previous lick. The code for lickometers and data from lickometry is available at https://github.com/dlhagger.

## Statistical analysis

Statistical analysis was performed with GraphPad Prism software (v. 9.0; GraphPad Software, La Jolla, CA) for all experiments except for lickometry datasets. For lickometry data, statsmodels, scipy, and pingouin were used to perform statistical analyses. Significance of data with multiple groups was evaluated via two-tailed one-way or two-way ANOVA with Tukey’s post hoc test dependent upon whether there were two types of variables (i.e. treatment group and mouse genotype) or a single variable (i.e. different treatment groups). For comparisons with three or more variables (i.e. genotype, gender, time), mixed-ANOVAs or mixed-linear models with the variables assigned as fixed effects, and all other variables present in the data set as random effects was used. Lickometry datasets were assessed using mixed-ANOVAs or mixed-linear models with the variables assigned as fixed effects, and all other variables present in the data set as random effects. Two sample continuous distributions were compared using two-sample Kolmogorov-Smirnov tests. Comparisons were considered statistically significant at P values of < 0.05.

## Results

### ADH1 deficiency or inhibition decreases ethanol consumption

To determine how peripheral ethanol metabolism affects ethanol consumption, we tested *Adh1* KO mice in the DID and 2BC paradigms of ethanol consumption. Both male and female *Adh1* KO mice consumed less ethanol in the binge-like DID paradigm than their wild-type (WT) counterparts across the experiment **(Fig. 2A)**. The mice were then subjected to the 2BC paradigm of drinking behavior where both male and female *Adh1* KO mice showed decreased ethanol preference and consumption **(Fig. 2B)**. A similar pattern of 2BC consumption was seen in mice not pre-exposed to drinking, indicating that the decreased preference and ethanol consumption exhibited by *Adh1* KO mice especially at higher ethanol concentrations was not driven by previous exposure to 20% ethanol **(Fig. S1)**. In addition, CYP2E1 and ALDH2 protein expression and ALDH2 activity were not changed in *Adh1* KO mice **(Fig. S2)**.

**Figure 2:**
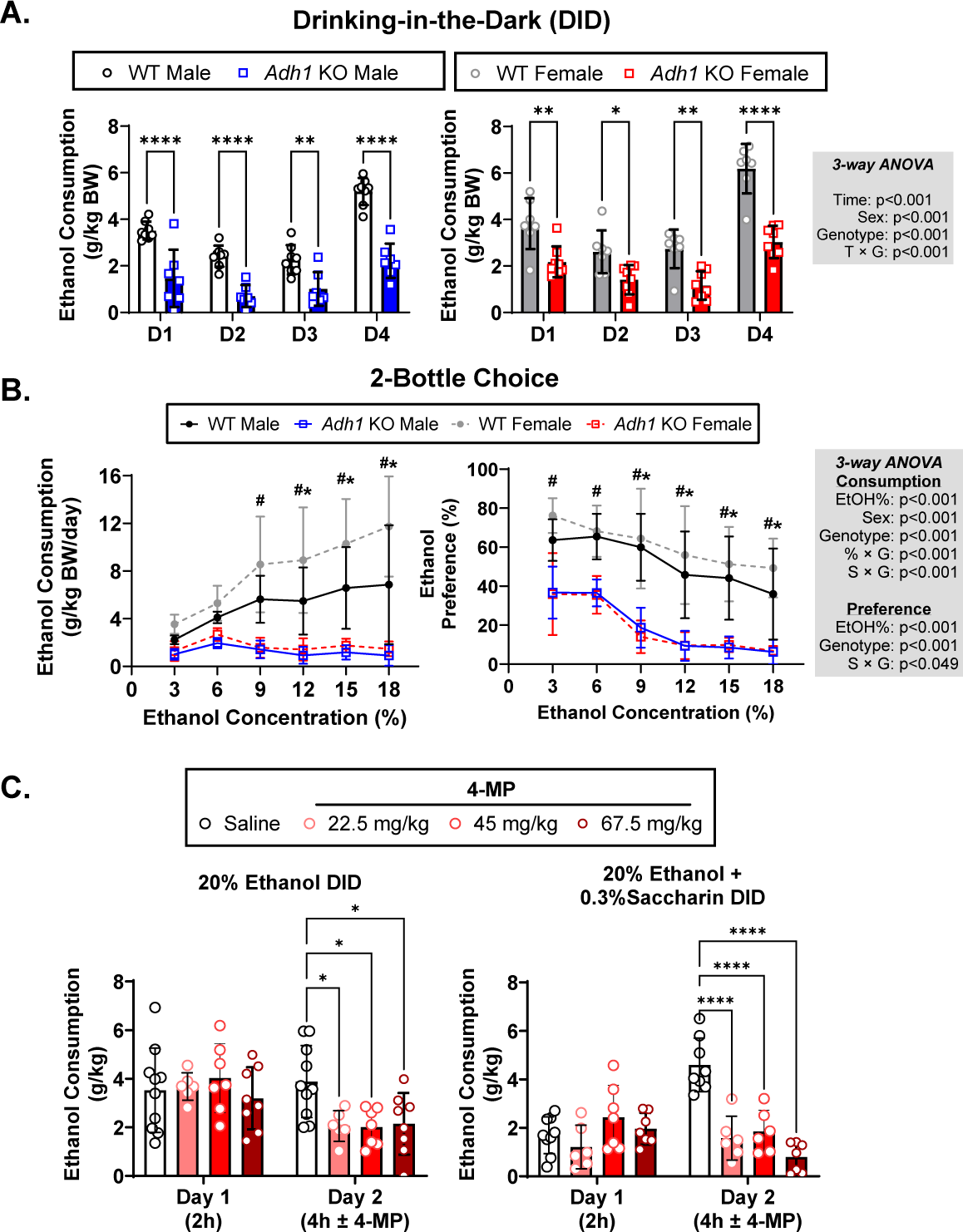
Deficiency or inhibition of ADH1 reduces ethanol consumption and preference. Age-matched male and female WT (C57BL6/N) or *Adh1* KO mice were first subjected to the drinking-in-the-dark (DID) paradigm (A) and then subsequently subjected to the 2-bottle choice paradigm (B) of drinking behavior. Female C57BL6/N mice (∼4mo) were subjected to the modified DID experiment evaluating consumption of normal or sweetened 20% ethanol (C). Baseline drinking values were collected on Day 1 (D1, 2h) before randomizing mice to groups for treatment on Day 2 (D2, 4h) with saline or different doses of ADH1 inhibitor fomepiezole (4-MP) (C). Values represent the mean ± SD (n=6-8 mice per group). Multiple-way ANOVA was used to analyze the significance of variable interactions with Tukey post-hoc test to identify differences between specific groups. For panels A and C \**P*<0.05, \*\**P*<0.01, \*\*\**P*<0.001, \*\*\*\**P*<0.0001 as indicated. For panel B, genotype comparisons at each ethanol concentration were assessed with ^#^*P*<0.05 for female mice and \**P*<0.05 for male mice.

While previous studies have shown that ADH1 inhibitor 4-MP can inhibit ethanol consumption in ethanol-preferring rats at a dose of 66.5 mg/kg via intraperitoneal (I.P.) injection, it is possible that 4-MP was exhibiting off-target effects (MacDonald, 1976). Since *Adh1* exhibits significant expression in the gastrointestinal tract, we hypothesized that we could give 4-MP via oral gavage which produces similar systemic 4-MP levels as intravenous injection (Marraffa et al., 2008), at reduced dosages to reduce potential off-target effects while maintaining ADH1 inhibition. We used C57BL6/N mice in a modified DID paradigm where mice were acclimated to ethanol consumption on day 1 (2h) and then treated with saline or 4-MP at different doses on day 2 (4h) with normal or sweetened 20% ethanol. Treatment with 4-MP reduced normal and sweetened ethanol consumption at all concentrations, indicating that ADH1 inhibition can also decrease ethanol consumption **(Fig. 2C)**.

### Ethanol and metabolite distributions are altered in *Adh1* KO mice

We then sought to determine how the lack of ADH1-mediated ethanol metabolism modulates the distribution of ethanol, AcH, and acetate in the blood, liver, and brain post ethanol gavage (2 g/kg). Ethanol levels in *Adh1* KO mice were not significantly different after 15 min, but were elevated in the blood, liver, or cerebellar cortex at the 1h timepoint **(Fig. 3A)**. AcH levels were decreased in the blood and liver at 15min and in the liver at 1h in *Adh1* KO mice compared to WT, while there was no difference in cerebellar AcH **(Fig. 3B)**. However, ethanol-induced increases in acetate were completely abolished in *Adh1* KO mice in serum, liver, and cerebellar cortex at all time points compared to WT mice after ethanol gavage **(Fig. 4A)** or I.P. injection **(Fig. 4B)**.

**Figure 3:**
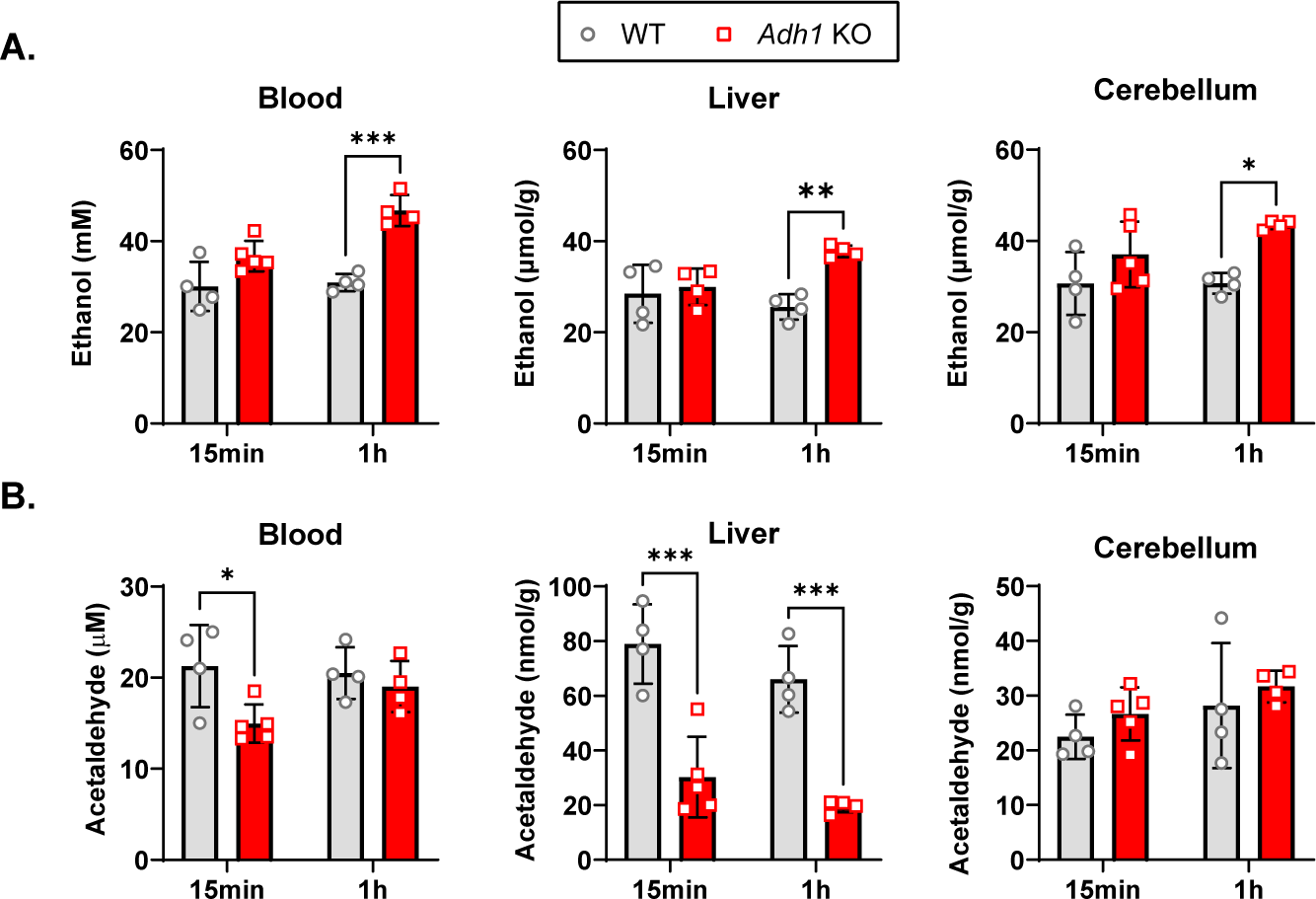
*Adh1* KO mice exhibit higher ethanol and lower peripheral acetaldehyde levels after ethanol administration. WT (C57BL6/N) or *Adh1* KO male mice (∼4mo) were given a 2g/kg gavage and sacrificed either 15min or 1h afterwards for collection of blood/serum, liver tissue, and cerebellar cortex. (A) Ethanol and (B) acetaldehyde (AcH) were measured via headspace GC-MS in blood, liver, and cerebellar cortex. Values represent the mean ± SD (n=4-5 mice per group). Two-way ANOVA was used to analyze the significance of variable interactions with Tukey post-hoc test to identify differences between specific groups.

**Figure 4:**
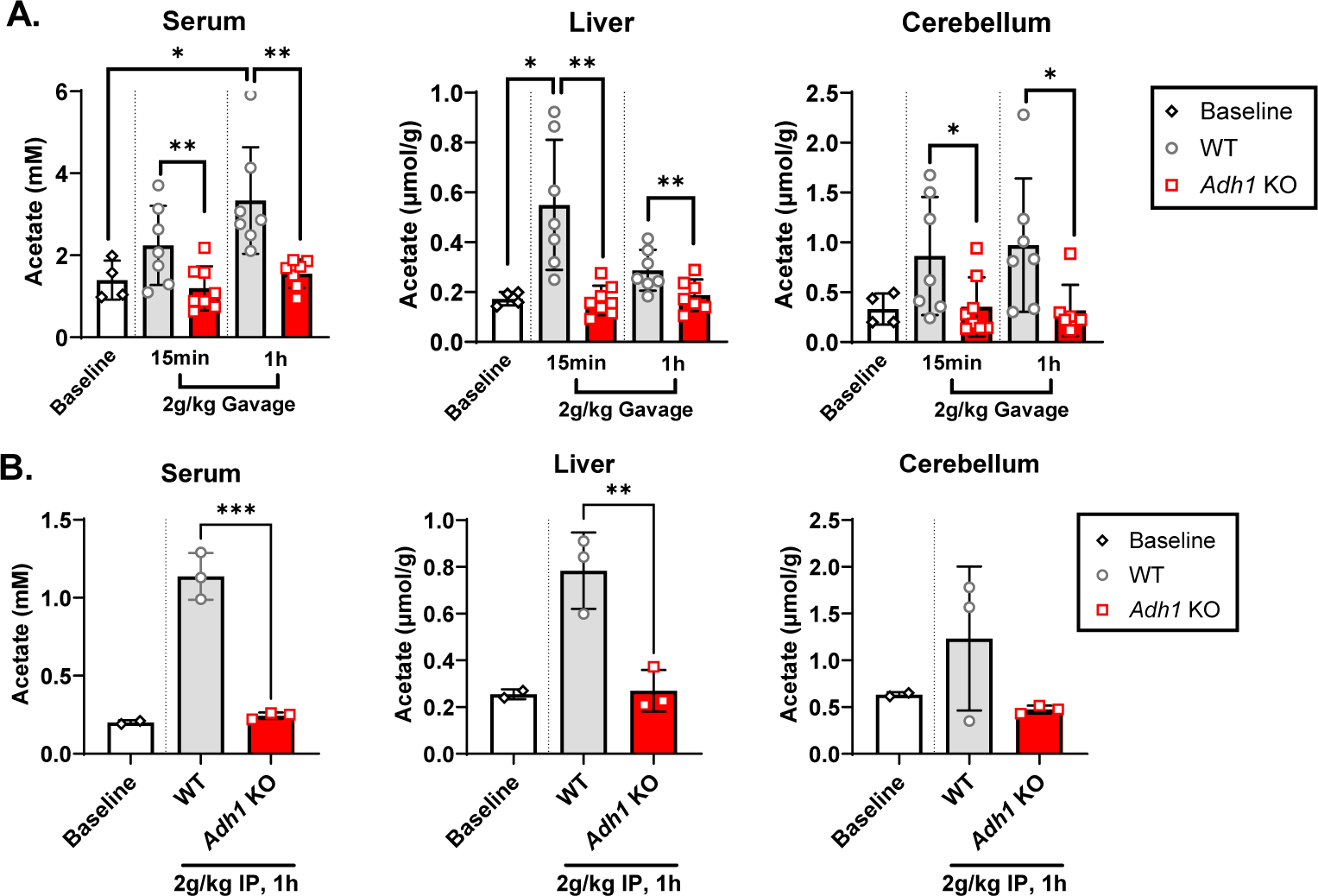
*Adh1* KO mice exhibit reduced acetate levels after ethanol administration. WT (C57BL6/N) or *Adh1* KO male mice (∼4mo) were given a 2g/kg gavage (A) or intraperitoneal injection (B) and sacrificed either 15min or 1h afterwards for collection of blood/serum, liver tissue, and cerebellar cortex. Acetate was measured via GC-MS in blood, liver, and cerebellar cortex. Values represent the mean ± SD (n=4-8 mice per group for (A) and n=2-3 mice per group for (B)). Two-way ANOVA was used to analyze the significance of variable interactions with Tukey post-hoc test to identify differences between specific groups.

### Ethanol levels are increased after voluntary consumption in *Adh1* KO mice

It was surprising to us that ethanol levels were not significantly different at early time points in *Adh1* KO mice. While ethanol gavage is the closest administration method to ingestion, there are still significant differences between voluntary ethanol consumption and administration via gavage. Therefore, we decided to measure ethanol levels in WT and *Adh1* KO mice after voluntary ethanol consumption during DID. While *Adh1* KO mice consumed about ∼50% of the ethanol compared to WT mice, they exhibited similar or higher levels of ethanol in the blood, liver, and brain at 2hr **(Fig. 5A-B)**. Analyzing the correlation between blood ethanol levels and consumption, we found that *Adh1* KO mice accumulate ethanol at 2-fold the rate of WT mice after DID ethanol consumption **(Fig. 5C)**. This was repeated in the modified DID paradigm after 4-MP treatment **(Fig. 5D)** and we found that even the lowest dose of 4-MP given orally was able to recapitulate the blood ethanol accumulation seen in *Adh1* KO mice **(Fig. 5E)**.

**Figure 5:**
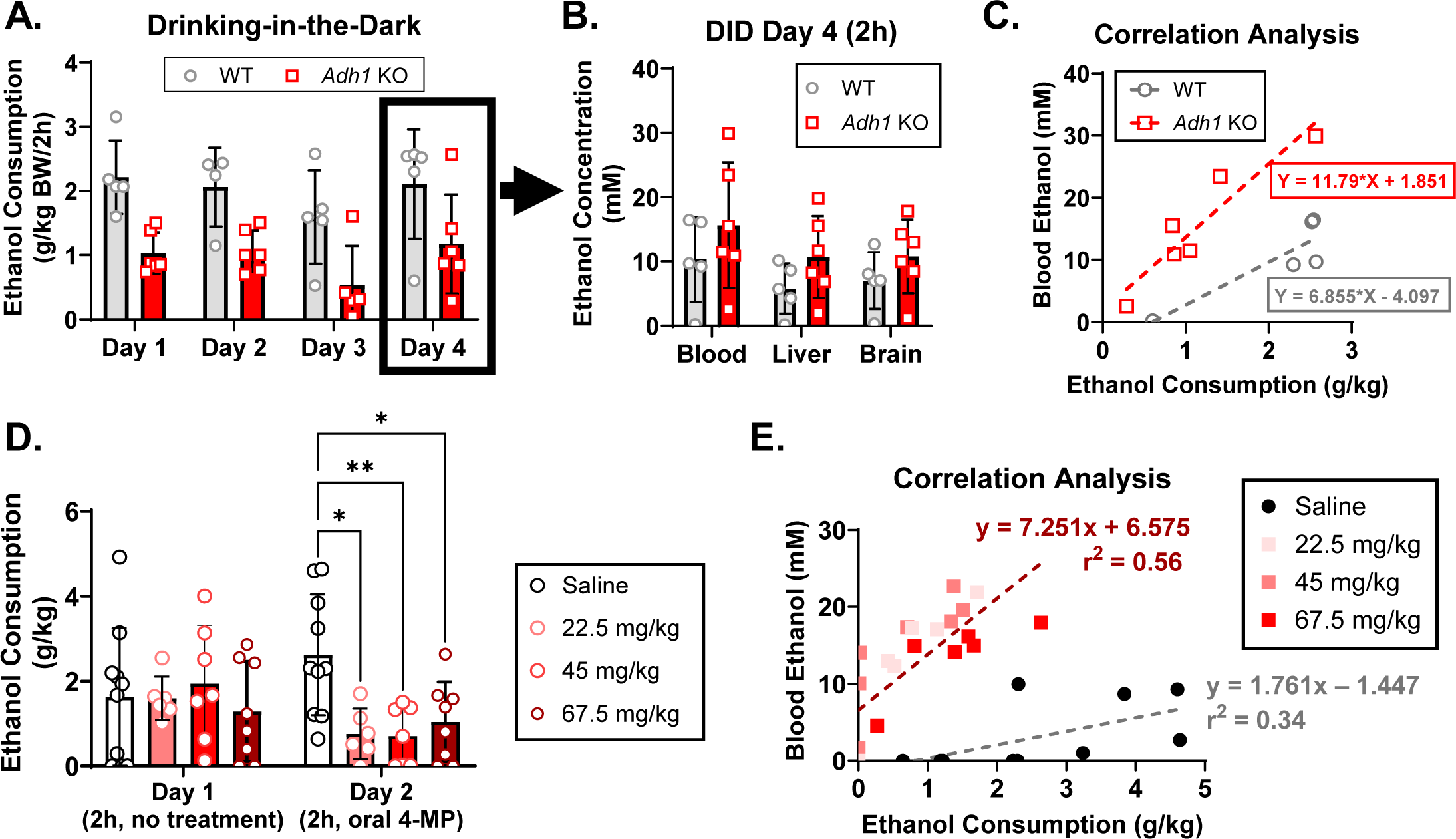
Ethanol levels in *Adh1* KO and 4-MP treated mice are significantly higher than control mice after DID. Age-matched male and female WT (C57BL6/N) or *Adh1* KO mice were subjected to the drinking-in-the-dark (DID) paradigm (A). On Day 4, mice were sacrificed after 2h of ethanol consumption for collection of blood/serum, liver tissue, and cerebellar cortex, and ethanol concentrations were measured via headspace GC-MS (B). Correlation analysis between blood ethanol levels and DID ethanol consumption (C). C57BL6/N mice from Fig. 1 were subjected to another round of modified DID (D). Blood was taken via orbital sinus 2h after the start of DID on Day 2 and measured. The blood ethanol level was plotted against measured ethanol consumption, and linear correlations for saline (grey) compared to all 4-MP treated mice (burgundy) were calculated (E). Values represent the mean ± SD (n=5-6 mice per group). Two-way ANOVA was used to analyze the significance of variable interactions with Tukey post-hoc test to identify differences between specific groups. \**P*<0.05, \*\**P*<0.01

### ADH1-mediated ethanol metabolism affects drinking microstructure

To dissect how ethanol consumption differs between WT and *Adh1* KO mice, we built a capacitance-based lickometry system based on previous studies (Petersen et al., 2023, Haggerty et al., 2022) and used it to study home-cage drinking behavior in WT and *Adh1* KO mice in the modified DID paradigm. In experiments containing both male and female WT and *Adh1* KO mice, we found that there were significant differences in the correlations between alcohol intake and total licks, total lick duration, and bouts, indicating vastly different drinking structures between the two genotypes used to achieve their associated alcohol intakes **(Fig. 6A-D)**. On the other hand, there were no differences in these correlations between WT and *Adh1* KO mice when consuming the noncaloric sweetener saccharin (0.1%) in water, indicating that the altered consumption behavior in *Adh1* KO mice is specific to alcohol **(Fig. 6E-H)**.

**Figure 6.**
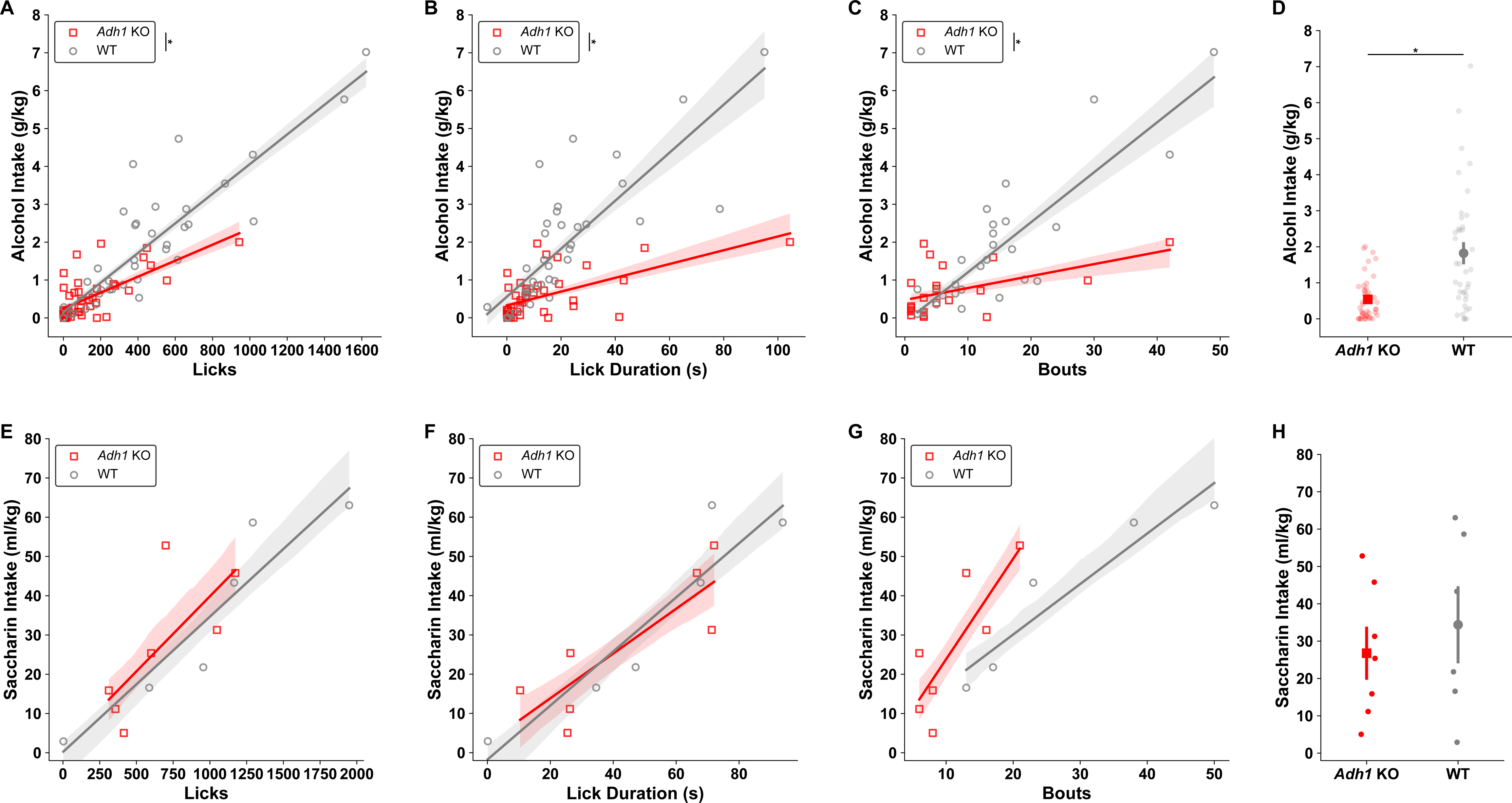
Drinking behaviors reliably predict DID intakes and alterations in *Adh1* KO drinking behaviors are alcohol-specific. *Adh1* KO mice drinking behaviors compared to wild-type mice predict decreased alcohol intakes **A)** as a function of decreased licks by genotype (MixedLM - Intake ∼ Licks * Genotype; Licks * Genotype *F_(1,64)_* = 22.76, *p* = 1.102e^-05^), **B)** as a function of decreased lick durations by genotype (MixedLM - Intake ∼ Lick Duration * Genotype * Sex; Lick Duration * Genotype *F_(1,64)_* = 48.59, *p* = 2.088e^-09^) and **C)** as a function of decreased bout numbers by genotype (MixedLM - Intake ∼ Bout Number * Genotype * Sex; Bout Number * Genotype *F_(1,45)_* = 21.44, *p* = 3.112e^-05^). **D)** Across all DID sessions where lickometry was measured, *Adh1* KO mice drank less alcohol than wild-type mice (MWU, *z* = 3.275, *p* = 0.001059). Whereas for saccharin drinking, *Adh1* KO mice drinking behaviors compared to wild-type mice do not differ by genotype, but do predict saccharin intake as a function of **E)** licks (MixedLM - Intake ∼ Licks * Genotype; Licks *F_(1,9)_* = 7.259, *p* = 0.02462), **F)** as a function of lick duration (MixedLM - Intake ∼ Lick Duration * Genotype; Lick Duration *F_(1,9)_* = 18.25, *p* = 0.002071), **G)** and as a function of bout numbers (MixedLM - Intake ∼ Bout Number * Genotype; Lick Duration *F_(1,8)_* = 12.96, *p* = 0.006969). **H)** Across all DID sessions where lickometry was measured, *Adh1* KO mice drank similar amounts of saccharin as wild-type mice (t-test, *t* = 0.6355, *p* = 0.5408). Individual data points are cumulative measurements from individual animals across DID sessions, shading and error bars represent SEM for lines of best fit and group means.

To further investigate how the drinking behavior differs between the two genotypes, we investigated how the licks differed over time for each genotype. Individual traces for cumulative licks across time in the DID session were plotted the by genotype for the 2h **(Fig. 7A)** DID sessions. In **(Fig. 7B)**, cumulative licks are plotted individually to visualize exactly when, and not, animals are drinking by genotype. Kernel density estimates from all x values (the time at which those licks occurred) and y values (how many licks have occurred) were computed and compared. *Adh1* KO mice have less licks for alcohol across time, and when these licks occur differs greatly compared to wild-type mice. Since the times at which mice licked differed by genotype, we further assessed frontloading behaviors by genotype. Although there is a visual difference showing less cumulative licks compared to wild-type mice during the frontloading period, this was not statistically significant **(Fig. 7C)**. On the other hand, *Adh1* KO mice have decreased numbers of cumulative licks post-frontloading **(Fig. 7D)**. The 4h DID sessions exhibit similar changes to the 2h DID in the cumulative lick and time domains, as well as front loading behavior **(Fig. 7E-G)**. Yet, post-frontloading, there is no difference by genotype **(Fig. 7H).** We believe this is due to the large density of drinking behavior that is observed between 120 and 150 minutes in 4h DID sessions for *Adh1* KO mice, which is denser than the frontloading period **(Fig. 7F)**. We further examined genotype-dependent differences in in licking microstructure. The licks over time are plotted as kernel density estimations split by bout identity **(Fig. 8A-B).** While the number of licks per DID session differ within bouts, interestingly, non-bout licks are similar between *Adh1* KO and wild-type mice **(Fig. 8A-B).** As a percentage of total licks, the number of licks that occur within a bout is decreased for *Adh1* KO mice, and while *Adh1* KO mice have similar numbers of non-bout licks during DID sessions, the durations for which they drink are almost twice as long compared to wild-type mice, with no difference within bouts **(Fig. 8C-D)**. Finally, as expected, *Adh1* KO mice have less bouts per session, lick fewer times within bouts, but do not have different total lick durations per bout, which also indicates a lower licking frequency per bout **(Fig. 8E-F).** These differences in drinking behavior are not likely to stem from worsened coordination in *Adh1* KO mice, as there were no differences accelerating rotarod performance between WT and *Adh1* KO mice after I.P. injection or gavage of ethanol **(Fig. S3)**.

**Figure 7.**
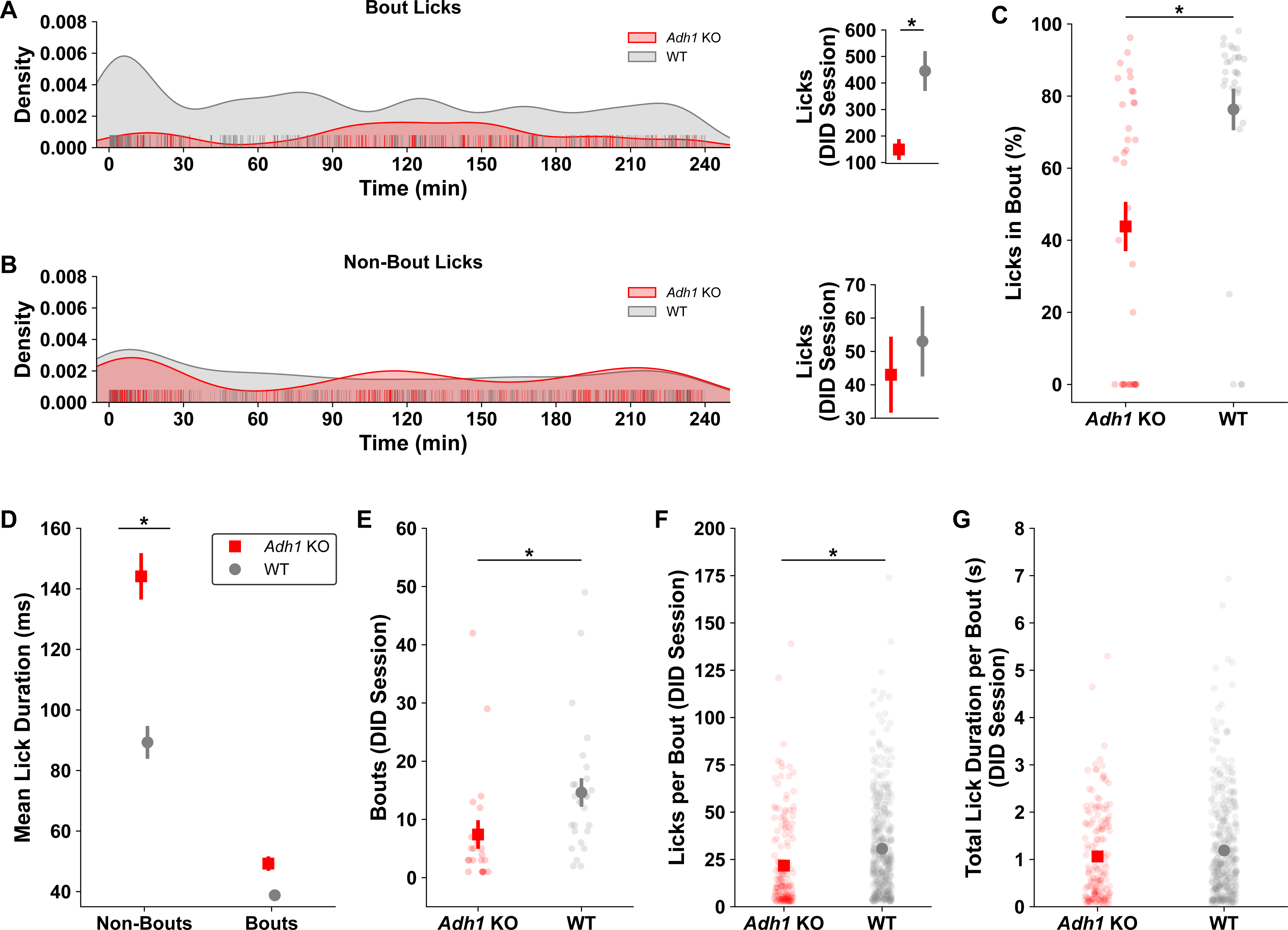
*Adh1* KO alcohol drinking behaviors display temporal shifts as a function of access to alcohol. **A)** Representations of *Adh1* KO and wild-type mice cumulative licks across time during a single two-hour DID session (lines represent individual animals). **B)** *Adh1* KO mice not only show difference in their cumulative licks compared to wild-type mice across the whole two-hour DID session (two-sample Kolmogorov-Smirnov – Cumulative Licks distributions by genotype; *KS* = 0.4232, *p* = 2.698e^-85^), but also show temporal differences in their drinking patterns across the two-hour DID session (two-sample Kolmogorov-Smirnov – Time (min) distributions by genotype; *KS* = 0.3944, *p* = 8.528e^-74^) (individual data points represent increases in cumulative lick measurements at the exact time they occurred for all animals). **C)** When assessing frontloading (first 30 min) behaviors by genotype, we failed to find differences in cumulative licks by genotype early in two-hour DID sessions (MWU, *z* = 1.610, *p* = 0.1179). **D)** Yet, when assessing drinking behaviors from 30 minutes to the end of the two-hour DID session, *Adh1* KO mice display decreases in licking behavior as compared to wild-type mice in this time window (t-test, *t* = 2.832, *p* = 0.02377). **E)** Representations of *Adh1* KO and wild-type mice cumulative licks across time during a single four-hour DID session (lines represent individual animals). **F)** Similar to two-hour DID sessions, *Adh1* KO mice not only show difference in their cumulative licks compared to wild-type mice across the whole four-hour DID session (two-sample Kolmogorov-Smirnov – Cumulative Licks distributions by genotype; *KS* = 0.2532, *p* = 2.134e^-93^), but also show temporal differences in their drinking patterns across the four-hour DID session (two-sample Kolmogorov-Smirnov – Time (min) distributions by genotype; *KS* = 0.1711, *p* = 1.164e^-42^) (individual data points represent increases in cumulative lick measurements at the exact time they occurred for all animals). **G)** When assessing frontloading (first 30 min) behaviors by genotype, we again failed to find differences in cumulative licks by genotype early in four-hour DID sessions (t-test, *t* = 1.826, *p* = 0.09691). Unlike the two-hour DID sessions though, when assessing drinking behaviors from 30 minutes to the end of the four-hour DID session, *Adh1* KO mice also show no difference in temporal licking behavior as compared to wild-type mice in this time window (MWU, *z* = 1.846, *p* = 0.06978). Individual data points are cumulative measurements from individual animals across DID sessions. Cumulative lick and time distributions are displayed as kernel density estimations from individual datapoints. Error bars represent SEM for group means.

**Figure 8.** Deficits in *Adh1* KO alcohol drinking microstructures drive decreases in alcohol intakes. Kernel density estimations of drinking behaviors across all DID sessions when split by drinking behaviors **A)** during bouts and **B)** outside of drinking bouts show that the total number of licks during bouts differs significantly by genotype (MWU, *z* = 3.457, *p* = 0.0005660), whereas the number of licks that occur during non-bouts is not different by genotype (MWU, *z* = 1.769, *p* = 0.07771). **C)** When sampling the licks based on their bout identity, *Adh1* KO mice also display a lower proportion of licks that occur in bouts than wild-type mice as a percentage of total licks (MWU, *z* = 4.099, *p* = 0.00003700). Finally, when looking the mean duration of licks both during bouts and non-bouts, we show that **D)** *Adh1* KO mice spend more time drinking alcohol during a single bottle contact that wild-type mice during non-bout licks, but no difference during bouts (mixed-ANOVA; Genotype * Bout Type *F_(1,18)_* = 4.954, *p* = 0.03903; Bouts contrast, *p* = 0.1738; Non-bouts contrast *p* = 0.04411). When looking at genotype differences only within bouts, as expected we find *Adh1* KO mice have **E)** less bouts (MWU, *z* = 3.386, *p* = 0.0007070), **F)** but also, fewer licks within those bout structures (MWU, *z* = 5.008, *p* = 5.403e^-13^), **G)** yet similar overall drinking times within those bout structures (MWU, *z* = 1.494, *p* = 0.1350).

## Discussion

Here, we show that peripheral ethanol metabolism, which is mainly mediated by ADH1, has a significant influence on ethanol consumption and is a potent modulator of drinking behavior in mice. ADH1-mediated changes in drinking behavior are likely due to changes in the central levels of ethanol and acetate as a function of their peripheral metabolism, as central AcH levels did not change between WT and *Adh1* KO mice.

These pharmacokinetic differences between WT and *Adh1* KO mice lead to vastly different drinking behaviors in time and in microstructures for ethanol but not saccharin, highlighting the important effect of peripheral ethanol metabolism in drinking behavior.

How ethanol levels themselves regulate ethanol consumption is controversial. One study showed that up to 10-fold increased ethanol levels via subcutaneous 4-MP treatment (81 mg/kg) did not alter 10% ethanol intake in mice in a 2BC consumption paradigm (Gentry et al., 1983). On the other hand, intravenous infusion of ethanol or treatment with 4-MP was able to decrease voluntary consumption of 10% ethanol (Waller et al., 1982), and 4-MP can decrease ethanol consumption in alcohol-preferring rats (Peana et al., 2017). Our DID data shows that oral administration of the lowest dose of 4-MP (22.5 mg/kg) exhibited ethanol accumulation and inhibition of drinking behavior similar to higher 4-MP doses and *Adh1* KO mice. Thus, the ability of 4-MP to alter drinking behavior seems to be dependent on dose, administration route, ethanol consumption paradigm, and the amount of ethanol consumed during the voluntary access period. Taken together, our data with both 4-MP and *Adh1* KO mice demonstrate that loss of ADH1 activity reduces binge-like and 2BC ethanol consumption in mice. While the exact understanding for how *Adh1* KO mice alter ethanol and its metabolites levels that lead to reduced ethanol consumption is yet to be fully elucidated, one observation is that mice may titrate their drinking to achieve blood ethanol levels that optimize for the positive effects of high ethanol levels, while minimizing the aversive effects. Our DID findings partially support this notion, as shifts in the temporal microstructures may be reflections of this phenomenon, but the reality is likely much more complex. Future studies of ethanol titration in this model will attempt to identify the contributions of peripheral metabolism to ethanol-specific tolerance and seeking.

There are two lines of evidence that suggest ADH-mediated peripheral ethanol metabolism is a key determinant of ethanol consumption levels in humans. The first is that genetic polymorphisms in *ADH1B* and *ADH1C* which increase the activity of ADH1 and the generation of AcH after ethanol consumption decrease the risk of AUD (Zhou et al., 2022, Kimura and Higuchi, 2011, Luczak et al., 2009). On the other hand, bariatric surgery, which reduces gastric first-pass ethanol metabolism and results in much higher levels of ethanol per drink after consumption, is a risk factor for AUD development (Blackburn et al., 2017, Acevedo et al., 2020, Woodard et al., 2011). Even though *Adh1* KO mice exhibit similar increases in ethanol levels after ethanol consumption as bariatric surgery patients due to the lack of first-pass ethanol metabolism, these mice consume significantly less ethanol. While this discrepancy is puzzling at first, bariatric surgery patients retain the ability to metabolize ethanol in the liver and other peripheral organs expressing ADH1. Therefore, bariatric surgery patients should be able to generate significant amounts of the ethanol metabolites AcH and acetate compared to our *Adh1* KO mice, which generate less AcH and little to no acetate from ethanol metabolism. In combination, the evidence suggests that ethanol metabolites, especially acetate, may play an important role in the acquisition and reinforcement of ethanol consumption.

With the recent evidence implicating exogenous (usually produced through fermentation of dietary fiber by microbiota) (Erny et al., 2021) and endogenous (pyruvate-derived) (Liu et al., 2018) acetate in organismal homeostasis, the lack of investigation into the function of acetate in ethanol-induced neuronal and behavioral changes is puzzling. For instance, acetate can improve metabolic fitness and cognitive function in preclinical models of sleep disruption by binding pyruvate carboxylase and restoring normal energy homeostasis (He et al., 2024). Acetate also modulates both acute and chronic feeding behaviors, brain innate immunity, and rescues social deficits in a preclinical autism spectrum disorder model (Frost et al., 2014, Perry et al., 2016, Erny et al., 2021, Osman et al., 2023). Ethanol-derived acetate plays important roles in ethanol-induced physiological changes through downstream conversion to GABA and brain histone acetylation, as well as NMDAR activation which mediates some ethanol effects on blood pressure and amygdala/accumbens shell neuron stimulation (Chapp et al., 2024, Chapp et al., 2023, Chapp et al., 2014, Mews et al., 2019, Jin et al., 2021). In our study, the most drastic metabolic change in *Adh1* KO mice was the complete ablation of ethanol-induced increases in acetate after ethanol administration. Though ethanol-derived acetate is important in ethanol-induced learning (Mews et al., 2019), the direct effects of acetate on ethanol consumption are unclear. Whether the altered alcohol consumption in *Adh1* KO mice involves the loss of acetate-mediated NMDAR activation or brain histone acetylation will be a topic of future study using acetate supplementation and NMDAR activators in *Adh1* KO mice. It is also important to note that the effects of decreased ALDH2 function via polymorphism or inhibition on drinking behavior have been mostly attributed to higher acetaldehyde levels; however, decreases in ALDH2 function also decrease ethanol-derived acetate levels (Kiyoshi et al., 2009), and the effects of this change on drinking behavior have not been rigorously investigated.

The intensity of frontloading behavior has been shown as an important predictor of subsequent ethanol consumption, especially in preclinical models, and is a potent mediator for increasing the risk for the development of AUD (Griffin III et al., 2009). However, there is still much work to be done to elucidate ethanol metabolism’s role in driving voluntary drinking and alcohol tolerance. Here, we described novel shifts in the temporal dynamics and structure of ethanol intake due to changes in peripheral metabolism alone that are driving processes largely thought to be cognitive, centrally driven actions mediated by salience and interoception. A recent brain microdialysis study in a free drinking rat found that brain acetate and ethanol levels rose at the same rate, indicating that ethanol-derived acetate quickly accumulates in the brain after voluntary ethanol consumption (Lee et al., 2024). These data, taken together with our measurement of ethanol and its metabolites across during DID could indicate that a lack of acetate may impair learned-alcohol behaviors that lead to drinking escalation when ethanol levels are in similar ranges across genotypes. Evidence from the 2-hour DID sessions, in which *Adh1* KO mice have no chronic alcohol history, also shows that temporal dynamics for and the bout structures of voluntary alcohol drinking is already different from wild-type controls within minutes of consumption, which indicates there may also be innate differences in drinking phenotypes due to genetic rather than pharmacological manipulations of the machinery that metabolizes alcohol. Together, these observations provide a platform upon which future studies leveraging direct genetic manipulations of alcohol metabolism can be used to elucidate how specific changes in the pharmacokinetics of ethanol metabolism can lead to changes in alcohol-related behavior.

While our study provides important information for the alcohol field, it does have limitations. ADH1 is an important enzyme in retinoic acid disposition, and the *Adh1* KO mice may develop differently than their WT counterparts. In addition, we did not use littermate controls for *Adh1* KO experiments to simplify animal breeding. However, we validated that our findings are related to ADH1 function using 4-MP in littermate control animals. We also evaluated lickometry only in animals pre-exposed to at least 1 DID session to ensure a more consistent alcohol intakes which obscures potential differences in acquisition of ethanol drinking during the first exposure.

Overall, ethanol metabolism is a complex process that dictates many of the functional and behavioral changes induced by ethanol. Our findings highlight the importance of ADH1-mediated peripheral ethanol metabolism in dictating the peripheral and central pharmacokinetics of ethanol and its metabolites, which clearly drives different ethanol consumption behaviors. Future studies will deepen our understanding of how these metabolic changes affect ethanol tolerance, ethanol interoception, and brain circuits and regions important for the acquisition and maintenance of ethanol consumption.

## Acknowledgements

This work was supported by the intramural program of NIAAA, NIH (BG, DML) and NIGMS, NIH 1FI2GM154674-01 (DLH).

## Conflicts of interest

All authors disclose no conflicts.

## Author Contributions

B.M. and D.L.H. designed and performed experiments, analyzed the data, and edited the manuscript. T.L., Y.L., Y.F., H.L., R.J.P., T.R., W.S., D.F. performed some experiments and data analysis, and edited the paper. L.Z. helped with data analysis, provided relevant intellectual input, and edited the manuscript. D.M.L. and B.G. obtained funding and provided supervision. B.M. supervised the whole project and wrote the paper. All authors approved the final manuscript

---

**Figure S1. *Adh1* KO mice exhibit similar drinking patterns in 2BC without previous DID.** Ethanol-naïve male WT and *Adh1* KO mice were subjected to 2BC paradigm at escalating ethanol concentrations (5, 10, 15, 20%).

**Figure S2. *Adh1* KO mice exhibit normal ALDH2 expression and activity.** Protein from livers of WT and *Adh1* KO mice after DID (from Fig. 4) were extracted. Western Blot detection of ALDH2 and CYP2E1 was performed with β-actin as the loading control (A). ALDH2 activity was assessed (B).

**Figure S3. *Adh1* KO mice do not exhibit more impairment than WT mice after low-dose ethanol administration at early timepoints.** Age-matched male WT and *Adh1* KO mice were subjected to accelerating rotarod after 1 g/kg (A) or 2 g/kg (B) intraperitoneal (I.P.) ethanol injections or a 2 g/kg ethanol gavage (C).

## Notes

### Competing Interest Statement

The authors have declared no competing interest.

