## Supplementary figures and images for "Peripheral alcohol metabolism dictates ethanol consumption and drinking microstructure in mice"

### Supplemental Figure 1

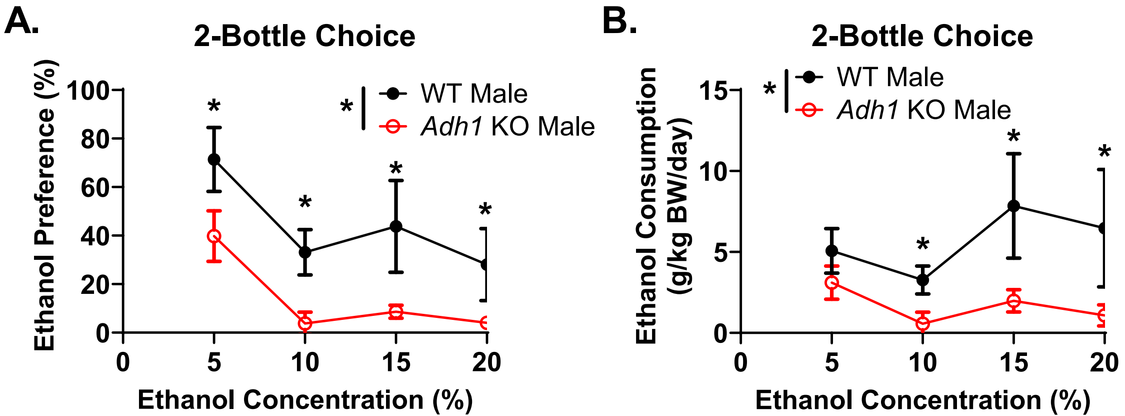

### Supplemental Figure 2

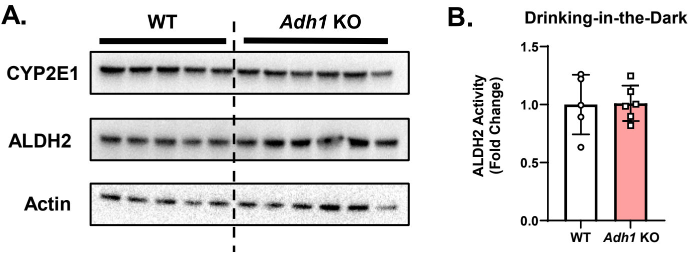

### Supplemental Figure 3

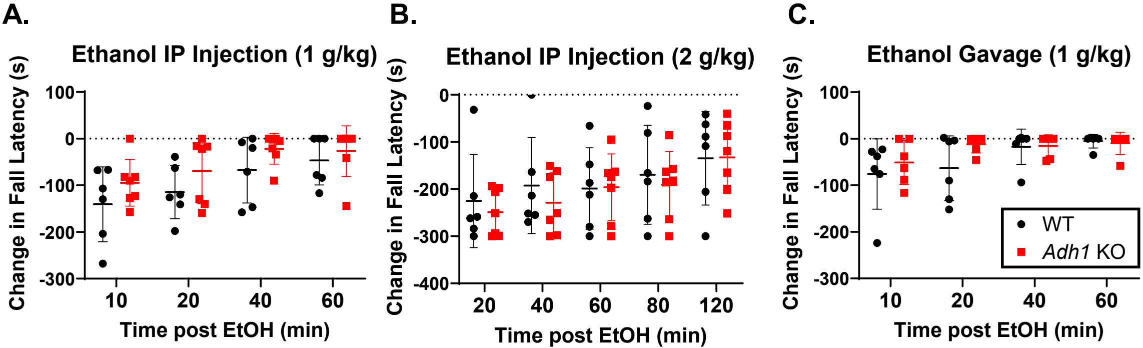
